# Anatomical determinants of DTI-ALPS: effects of ROI definition, ventricular morphology, and periventricular deformation

**DOI:** 10.64898/2026.07.02.735989

**Authors:** David K Wright

**Author notes:** **Corresponding author information:** David K Wright, Department of Neuroscience, The School of Translational Medicine, Monash University, Level 6, Alfred Centre, 99 Commercial Road, Melbourne, VIC 3004, Australia, E.

## Abstract

The diffusion tensor image analysis along the perivascular space (DTI-ALPS) index is increasingly used as a non-invasive MRI biomarker of glymphatic and perivascular function, yet the anatomical validity and measurement stability of the metric remain incompletely characterised. Using diffusion MRI data from 850 healthy young adults and 150 healthy ageing participants from the Human Connectome Project, I systematically evaluated the influence of region-of-interest (ROI) placement and ventricular anatomy on ALPS measurements. Reference ALPS implementations demonstrated substantial hemispheric variability, with a median left-right difference of 12.5% and marked asymmetry in the underlying numerator and denominator tensor components. A two-stage optimisation framework incorporating fibre-pool alignment, hemispheric symmetry, component stability, and directional purity identified anatomically improved ROI configurations that significantly increased fibre specificity and reduced measurement variability in independent validation cohorts. Despite these improvements, residual hemispheric asymmetry persisted, suggesting an intrinsic anatomical contribution to ALPS variability. In the healthy ageing cohort, ventricular volume emerged as the strongest predictor of ALPS, explaining substantially more variance than chronological age. Voxel-wise deformation-based morphometry demonstrated that lower ALPS values were associated with ventricular and periventricular expansion, while optimisation increased coupling between ALPS and ventricular anatomy. Collectively, these findings indicate that ALPS measurements are strongly influenced by ROI definition, ventricular morphology, and surrounding periventricular tissue architecture. Rather than functioning as a direct measure of glymphatic transport in isolation, ALPS appears to represent a composite anatomical diffusion biomarker shaped by both methodological implementation and underlying neuroanatomy. These results provide a framework for improving methodological standardisation and interpretation of ALPS measurements in future neuroimaging studies.

## 1. INTRODUCTION

The brain’s capacity to clear interstitial solutes via perivascular pathways has emerged as a key mechanism in maintaining homeostasis and is increasingly implicated in aging and neurodegenerative disease (Han et al., 2023; Harrison et al., 2020; Iliff et al., 2012; Ishida et al., 2022; Kress et al., 2014; Nedergaard & Goldman, 2020; Rasmussen et al., 2018; Ringstad et al., 2018; Zamani et al., 2022; Zamani et al., 2025). Consequently, considerable effort has been directed toward developing non-invasive imaging markers capable of interrogating these systems *in vivo* (Han et al., 2023; Harrison et al., 2018; Ohene et al., 2019; Ringstad et al., 2018; Taoka & Naganawa, 2020). Among the proposed approaches, diffusion tensor imaging analysis along the perivascular space (DTI-ALPS) has gained widespread adoption because it can be derived from conventional diffusion MRI acquisitions and has been reported to detect alterations across a broad range of neurological disorders and ageing populations (Han et al., 2023; S. Liu et al., 2024; Satpathi et al., 2025; Sharkey et al., 2024; Taoka et al., 2022; Taoka et al., 2024; Taoka et al., 2017; Yu et al., 2026).

DTI-ALPS is measured at the level of the upper lateral ventricle, where medullary veins run predominantly in the left-right direction and are approximately orthogonal to both projection and association fibre bundles (Taoka et al., 2022; Taoka et al., 2024). The method exploits this ordered anatomy by comparing diffusivity parallel and perpendicular to the expected orientation of projection and association fibres, generating the ALPS index as a surrogate measure of perivascular diffusivity.

Despite its rapid adoption, remarkably little work has examined the methodological behaviour of ALPS itself (Ringstad, 2024; Taoka et al., 2022). DTI-ALPS relies on placement of small regions of interest (ROIs) in projection and association fibre regions adjacent to the lateral ventricles, however there is no consensus regarding ROI size, shape, placement strategy or anatomical definition. Most studies have applied published ROI definitions directly to disease cohorts, implicitly assuming ALPS provides a stable and anatomically robust measurement. However, even small deviations in ROI placement can significantly alter the ALPS index (Taoka et al., 2024), particularly given the complex microstructural organisation of white matter in these regions and the spatial scale and variable orientation of perivascular spaces (Schilling et al., 2025).

Recent studies have raised broader questions regarding the biological specificity of ALPS measurements. Schilling and colleagues demonstrated that ALPS is influenced by white matter geometry, fibre crossings, and tensor orientation properties, suggesting that tissue microstructure may contribute substantially to the metric (Schilling et al., 2025). Similarly, Taoka and colleagues have emphasised that ALPS should be interpreted as a measure of directional water diffusivity within a highly specific anatomical region rather than as a direct measure of glymphatic transport (Taoka et al., 2024). Together, these observations suggest that both methodological implementation and underlying anatomy may exert important influences on ALPS measurements that remain poorly characterised.

In this study, I systematically evaluate the anatomical determinants of DTI-ALPS using large-scale diffusion MRI datasets from the Human Connectome Project Young Adult (HCP-YA) and Healthy Aging (HCP-HA) cohorts (Van Essen et al., 2011; Van Essen, Ugurbil, et al., 2012). A multi-stage optimisation framework incorporating fibre-pool alignment, hemispheric symmetry, component stability, and directional purity was developed to identify anatomically robust ROI configurations and assess the extent to which ROI placement contributes to measurement variability. In addition, relationships between ALPS, ventricular morphology, and periventricular tissue deformation were investigated using volumetric and deformation-based morphometry (DBM) analyses. Through this approach, the study aims to (i) quantify the influence of ROI definition on ALPS measurements, (ii) determine whether optimisation improves anatomical validity and measurement stability, and (iii) establish the extent to which ventricular anatomy contributes to variation in ALPS. These findings provide new insight into the interpretation, standardisation, and biological specificity of one of the most widely used diffusion MRI biomarkers of putative perivascular function.

## 2. METHODS

### 2.1 Ethics

This study was approved by the Monash University Human Research Ethics Committee (51994) and complied with the National Statement on Ethical Conduct in Human Research (2025).

### 2.2 Diffusion MRI dataset

Diffusion-weighted MRI data were obtained from two independent publicly available cohorts: the HCP-YA and HCP-HA cohorts (Table 1) (Van Essen et al., 2011; Van Essen, Ugurbil, et al., 2012). Both cohorts were analysed independently unless otherwise stated.

**Table 1.** Study datasets used for ROI optimisation, validation, and anatomical analyses. The Human Connectome Project Young Adult (HCP-YA) cohort was divided into a 50-subject development dataset used for ROI optimisation and an independent 800-subject evaluation dataset used for out-of-sample validation. Results are also presented for the combined HCP-YA cohort (n = 850). External validation of ventricular and deformation-based analyses was performed in 150 participants from the Human Connectome Project Healthy Aging (HCP-HA) cohort.

| Stage | Dataset | N |
| --- | --- | --- |
| Development | HCP-YA | 50 |
| Evaluation | HCP-YA | 800 |
| Combined | HCP-YA | 850 |
| Validation | HCP-HA | 150 |

The HCP-YA cohort comprised 850 neurologically healthy young adults with complete minimally pre-processed diffusion MRI datasets (Andersson et al., 2003; Andersson & Sotiropoulos, 2015; Andersson & Sotiropoulos, 2016; Glasser et al., 2013) acquired using multiband EPI (Feinberg et al., 2010; Moeller et al., 2010; Setsompop et al., 2012; Xu et al., 2012) at 3T. To facilitate unbiased optimisation and validation, an initial development subset of 50 subjects was randomly selected for ROI optimisation and methodological refinement, while the remaining 800 subjects were reserved for independent out-of-sample evaluation.

External validation was performed using 150 participants from the HCP-HA cohort spanning middle and older adulthood. This cohort was selected to evaluate the generalisability of the optimisation framework in the context of age-related ventricular enlargement and structural brain changes not represented in the young adult cohort.

HCP-HA subjects were selected using ventricular-volume stratified sampling across quintiles, with deliberate retention of ventricular extremes. This strategy was chosen because ventricular volume was strongly associated with age (r = 0.67) but remained heterogeneous after accounting for age and sex (R² = 0.49), enabling investigation of ventricular anatomy beyond chronological aging.

In addition to diffusion MRI, high-resolution T1-weighted structural images and pre-processed data (Fischl, 2012; Jenkinson et al., 2002; Milchenko & Marcus, 2013; Van Essen, Glasser, et al., 2012) including associated non-linear transformation files were obtained for all HCP-HA participants. These images were used to derive deformation-based morphometry measures, including voxel-wise log-Jacobian maps and ventricular volume estimates, enabling investigation of relationships between ALPS metrics, ventricular enlargement and periventricular tissue deformation.

### 2.3 Diffusion MRI processing and ALPS quantification

Pre-processed diffusion-weighted MRI data from the HCP-YA and HCP-HA cohorts were processed using a tensor-based pipeline restricted to b=0 and b=1,000 s/mm² shells, and b=0 and b=1,500 s/mm² shells, respectively. Diffusion tensors were fit using FSL (dtifit) (Jenkinson et al., 2012; Smith et al., 2004), and directional diffusivities (Dxx, Dyy, Dzz) were computed in native image space.

The DTI-ALPS index was calculated using established directional diffusivity relationships within periventricular projection and association fibre regions (Taoka et al., 2017). For each hemisphere:

- Numerator: mean diffusivity along the x-axis (Dxx) sampled in projection and association ROIs
- Denominator: mean diffusivity perpendicular to projection fibres (Dyy) and association fibres (Dzz)

The hemispheric ALPS index was calculated as:

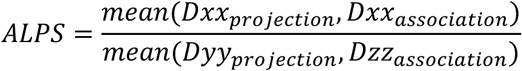

Bilateral ALPS was defined as the mean of left and right hemisphere values. Hemispheric asymmetry was quantified as the percentage difference between left and right ALPS values:

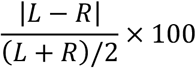

To identify sources of instability, numerator and denominator components were analysed separately using analogous left–right asymmetry metrics.

### 2.4 Reference ROI and fibre-pool generation

Various MNI coordinates, ROI sizes and ROI geometries have been used to define DTI-ALPS ROIs (Ayral et al., 2025; X. Liu et al., 2024; Qiu et al., 2025; Roura et al., 2025; Sharkey et al., 2024; Yun et al., 2025). In the present study, the term *reference ROI* refers to the literature-derived coordinate set adopted as the baseline implementation for optimisation and validation analyses. Reference DTI-ALPS ROIs were defined in MNI space at coordinates adjacent to the lateral ventricles within projection fibre regions (24, −12, 24 and −28, −12, 24) and association fibre regions (36, −12, 24 and −40, −12, 24) (Han et al., 2023; Yu et al., 2026). Small spherical ROIs approximating a 5 mm diameter were centred within white matter regions corresponding to the corticospinal tract and superior longitudinal fasciculus. Reference ROI masks were generated in MNI space and subsequently transformed into native diffusion space for each subject using spatial normalisation transforms provided by the HCP preprocessing pipelines. (Glasser et al., 2013), where subject-specific spherical ROIs were reconstructed prior to metric extraction.

To characterise the anatomical solution space available for optimisation, directional fibre pools were generated from the HCP-YA development cohort (n = 50) and mapped into MNI space. Projection-like and association-like white matter voxels were identified using directional dominance criteria applied to Dxx, Dyy and Dzz maps, and aggregated to form group-level projection and association reference pools. These anatomically constrained fibre pools served as reference regions for subsequent optimisation procedures. Reference ROIs and fibre pools were transformed between template and native subject diffusion space using the spatial normalisation transforms provided by the HCP preprocessing pipelines, enabling direct comparison between reference-defined ROIs and optimised subject-level solutions while preserving anatomical correspondence across participants.

### 2.5 Multi-criteria ROI optimisation framework

ROI optimisation was performed independently within the HCP-YA and HCP-HA cohorts using an identical two-stage framework designed to separate anatomical alignment from statistical refinement. Optimisation was initially developed using the HCP-YA development subset (n = 50), subsequently validated in the independent HCP-YA evaluation cohort (n = 800), and then repeated within the HCP-HA cohort (n = 150) to assess whether similar anatomically optimal solutions emerged in an independent ageing population.

### 2.6 Anatomical overlap optimisation

In the first stage, reference association ROIs were systematically shifted within a broad spatial search window (±4 mm in 1 mm increments) surrounding the published coordinates. Candidate locations were ranked according to overlap with the independently derived association fibre pool. The optimal left and right association ROI centres were identified at (38, −11, 27) mm and (−38, −13, 27) mm in MNI space, respectively, and were used as the anatomical starting point for subsequent multi-criteria optimisation.

### 2.7 Local refinement and candidate evaluation

In the second stage, local optimisation was performed within a restricted neighbourhood (±2 mm in 1 mm steps) surrounding the anatomically aligned ROI centres. Candidate ROIs were evaluated using a multi-criteria framework incorporating:

- hemispheric ALPS asymmetry,
- left–right ALPS bias,
- numerator and denominator stability,
- fibre purity and anatomical plausibility.

Candidate solutions were filtered using predefined anatomical and stability constraints, including minimum fibre overlap thresholds and maximum allowable hemispheric asymmetry.

Remaining candidate ROIs were ranked using a weighted composite scoring framework incorporating:

- hemispheric asymmetry (35%),
- ALPS bias (25%),
- component stability (40%).

This two-stage strategy explicitly decouples anatomical validity from statistical optimisation, reducing the risk of selecting ROIs solely based on favourable ALPS behaviour while ensuring structural plausibility of the final solution.

Reference, pool-optimised and multi-criteria optimised ALPS metrics were subsequently derived independently in the HCP-HA cohort using the same optimisation framework.

### 2.8 Hemispheric asymmetry and component analysis

To characterise measurement stability, hemispheric asymmetry analyses were performed for reference and optimised ALPS measurements across all subjects.

Absolute left–right percentage differences were calculated for:

- ALPS index,
- numerator terms,
- denominator terms.

Bland–Altman analyses were used to assess systematic hemispheric bias and limits of agreement between left and right ALPS measurements.

To determine whether apparent bilateral ALPS agreement masked instability in underlying tensor components, numerator and denominator asymmetry distributions were compared directly with overall ALPS asymmetry.

### 2.9 Out-of-sample validation

To assess generalisability of the optimisation framework, the final optimised ROI solution derived from the HCP-YA development subset was applied without modification to the independent HCP-YA evaluation cohort (n = 800). No re-optimisation was performed during this validation stage. Hemispheric asymmetry distributions obtained using reference and optimised ROIs were compared across the development subset, evaluation subset and combined full cohort to determine whether optimisation-derived improvements generalised beyond the original training sample.

To further assess robustness, the optimisation framework was independently applied to the HCP-HA cohort (n = 150). Anatomical overlap optimisation and subsequent multi-criteria optimisation were repeated using HCP-HA data, allowing direct comparison of reference, pool-optimised and multi-criteria optimised ALPS measurements within an independent ageing population.

### 2.10 Direction purity

Directional purity was quantified as the ratio of diffusivity along the expected fibre orientation to the mean diffusivity along the two competing orthogonal directions. Values >1 indicate increasing dominance of the intended fibre population, with larger values reflecting greater directional specificity of the ROI.

### 2.11 Ventricular volume and deformation-based morphometry

To investigate anatomical correlates of DTI-ALPS variability, ventricular volume and DBM analyses were performed in the HCP-HA cohort.

Lateral ventricular volumes were derived from the HCP structural processing pipeline and summed across hemispheres to obtain total lateral ventricular volume for each participant. Because ventricular volume exhibited a positively skewed distribution, log_10_-transformed ventricular volume was used for regression and visualisation analyses where appropriate. Voxel-wise deformation measures were obtained from the non-linear transformations generated during structural image registration to standard space (Fischl, 2012; Glasser et al., 2013; Jenkinson et al., 2002; Milchenko & Marcus, 2013; Van Essen, Glasser, et al., 2012). Log-Jacobian determinant maps were calculated from these deformation fields, providing a measure of local tissue expansion or contraction relative to the population template.

Whole-brain voxel-wise analyses were performed using permutation-based non-parametric inference implemented in FSL randomise (Winkler et al., 2014). Separate general linear models were constructed for reference and multi-optimised bilateral ALPS indices. Age and sex were included as nuisance covariates. Statistical significance was assessed using Threshold-Free Cluster Enhancement (TFCE) (Smith & Nichols, 2009) with family-wise error (FWE) correction. Results were considered significant at FWE-corrected *p* < 0.05.

To quantify local periventricular deformation, concentric ventricular shell masks were generated by morphological dilation of the lateral ventricle segmentation. Mean log-Jacobian values were extracted from 2-, 5-, and 10-voxel periventricular shells. Based on the voxel-wise DBM results and anatomical specificity considerations, the 2-voxel shell was selected *a priori* for subsequent quantitative analyses. Associations between ALPS metrics, ventricular volume, and periventricular log-Jacobian measures were assessed using linear regression.

### 2.12 Statistical analysis

Descriptive statistics are reported as mean ± standard deviation (SD) or median and interquartile range (IQR), as appropriate. Hemispheric asymmetry was quantified as the absolute percentage difference between left and right measurements relative to their mean. Bland–Altman analyses were performed to assess hemispheric agreement and estimate mean bias and 95% limits of agreement.

For optimisation and perturbation analyses, summary statistics were calculated across subjects for each candidate ROI configuration. Distributional properties were visualised using histograms, violin plots, Bland–Altman analyses, and density-based solution-space representations. Intermediate optimisation stages were evaluated using paired comparisons between reference and optimised metrics within the development cohort. Changes in hemispheric asymmetry, component stability, and directional purity were assessed using two-sided paired t-tests, with Wilcoxon signed-rank tests performed as sensitivity analyses where appropriate. Effect sizes are reported as paired Cohen’s *d*_z_.

Associations between ALPS metrics and demographic or anatomical covariates were assessed in the HCP-HA cohort using multivariable linear regression models including age, sex, and lateral ventricular volume. Continuous predictors were standardised prior to modelling. Separate models were fitted for reference and multi-optimised ALPS metrics. Model comparison analyses included age-only, ventricular–volume-only, and combined age–sex–ventricular-volume models. Incremental contributions of age and ventricular volume were assessed using partial F-tests comparing nested models. Unique variance explained by individual predictors was estimated as the reduction in model R² following removal of that predictor from the full model.

To evaluate potential non-linear relationships between ventricular anatomy and ALPS metrics, linear models were compared with natural spline models (3 degrees of freedom) using likelihood ratio tests.

False discovery rate (FDR) correction was performed using the Benjamini–Hochberg procedure. For covariate analyses, correction was applied separately for each outcome measure within each cohort and optimisation method, controlling for multiple testing across model predictors. For non-linearity analyses, FDR correction was applied across tests within each model family.

Statistical significance was defined as *p* < 0.05.

## 3. RESULTS

### 3.1 Reference DTI-ALPS ROIs exhibit substantial hemispheric variability

Reference DTI-ALPS ROIs were positioned adjacent to the lateral ventricles within projection and association fibre regions using previously published coordinates (Figure 1A). Visual comparison with anatomically derived directional fibre pools demonstrated that the projection ROIs were well contained within the projection tract core, whereas the association ROIs partially extended beyond the highest-density association fibre regions.

**Figure 1.**
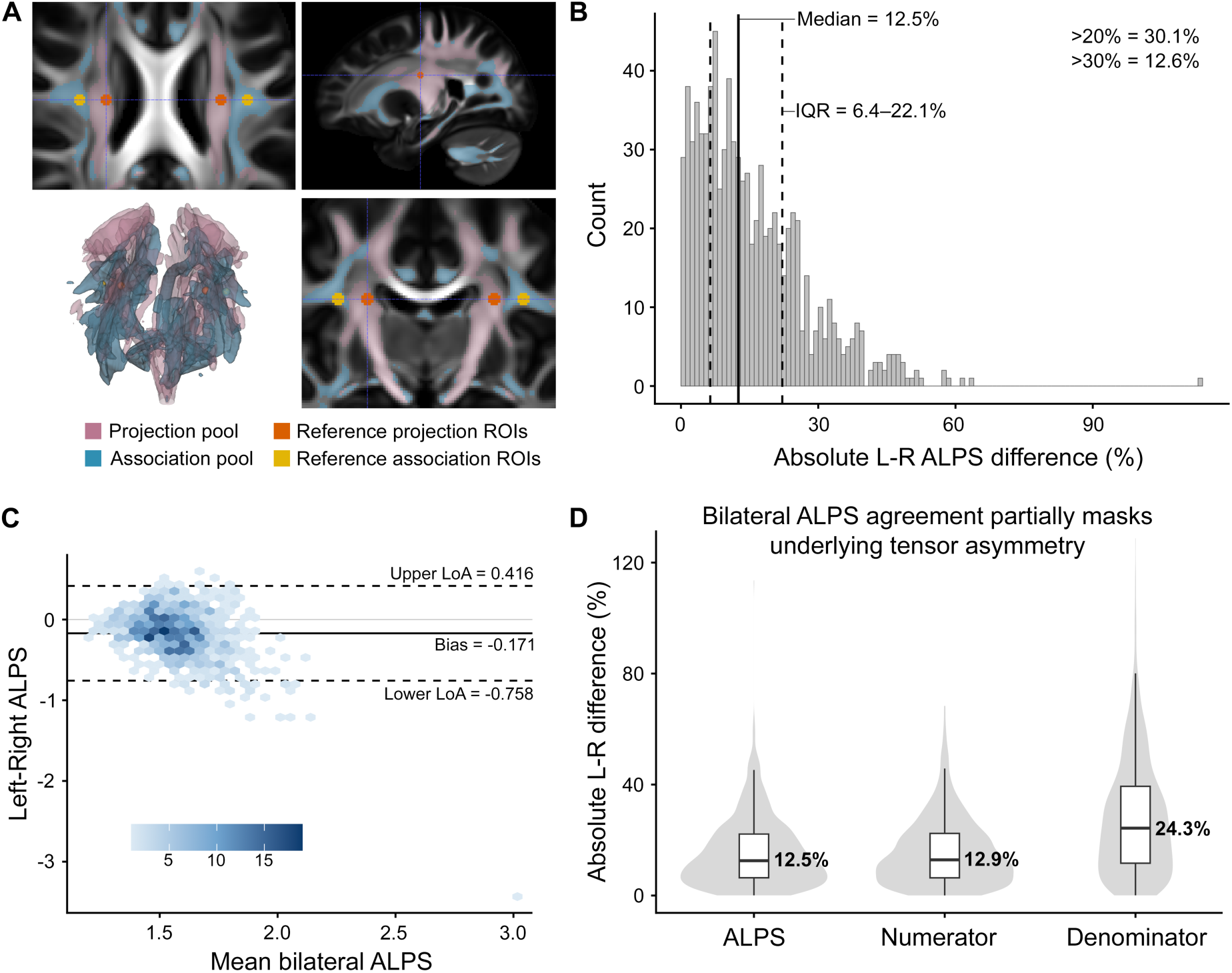
Reference DTI-ALPS regions of interest exhibit substantial hemispheric and component-level asymmetry. **(A)** Reference DTI-ALPS regions of interest (ROIs) overlaid on directional fibre pools derived from tensor-based tract orientation criteria. Projection ROIs (orange) are largely contained within the projection fibre pool (pink), whereas association ROIs (yellow) extend beyond the highest-density association fibre core (blue), suggesting imperfect anatomical alignment. **(B)** Distribution of absolute left–right (L–R) differences in the reference DTI-ALPS index across the full HCP-YA cohort (n = 850). The median hemispheric difference was 12.5% (IQR 6.4–22.1%), with 30.1% and 12.6% of participants exhibiting asymmetry greater than 20% and 30%, respectively. **(C)** Bland–Altman analysis of hemispheric agreement for reference DTI-ALPS measurements. A systematic rightward bias was observed (mean L–R difference = −0.171), with wide limits of agreement indicating substantial subject-level hemispheric variability. **(D)** Absolute hemispheric asymmetry distributions for the ALPS index and its constituent numerator and denominator terms. While median ALPS asymmetry was 12.5%, denominator asymmetry was markedly larger (24.3%), indicating substantial instability in the underlying tensor components. The lower asymmetry of the final ALPS ratio reflects partial cancellation of numerator and denominator biases rather than true hemispheric agreement.

Across the full HCP-YA cohort (n = 850), mean bilateral DTI-ALPS was 1.57 (left hemisphere: 1.49; right hemisphere: 1.66). Despite this apparently stable group-level estimate, substantial hemispheric variability was observed at the individual-subject level. The median absolute left–right difference in ALPS was 12.5% (IQR 6.4–22.1%), with 30.1% of participants exhibiting hemispheric differences exceeding 20% and 12.6% exceeding 30% (Figure 1B).

Bland–Altman analysis demonstrated a systematic hemispheric bias favouring higher right-hemisphere ALPS values (mean L–R difference = −0.171), with wide 95% limits of agreement ranging from −0.758 to 0.416 (Figure 1C). These findings indicate that reference ALPS measurements are not interchangeable between hemispheres and exhibit considerable subject-level variability despite relatively consistent cohort averages.

To determine whether bilateral ALPS agreement reflected stability of the underlying tensor components, numerator and denominator terms were analysed separately. Median absolute hemispheric differences were 12.9% for the numerator and 24.3% for the denominator, substantially exceeding the overall ALPS asymmetry (Figure 1D). The denominator component also demonstrated a strong signed hemispheric bias (median signed difference 20.1%), indicating that substantial asymmetry exists within the underlying diffusivity measurements and is partially obscured by cancellation effects within the ALPS ratio itself.

Collectively, these findings demonstrate that substantial instability exists within the constituent tensor components of the reference ALPS measurement, even in neurologically healthy young adults.

### 3.2 Association fibre-pool optimisation improves anatomical specificity but produces limited gains in ALPS stability

Association fibre-pool optimisation identified anatomically improved ROI locations centred at (38, −11, 27) mm and (−38, −13, 27) mm for the left and right association ROIs, respectively, corresponding to shifts of approximately 2–3 mm from the reference coordinates. Across the 50-subject development cohort, association pool optimisation produced a marked improvement in directional purity, increasing median association purity from 1.90 to 2.58 (paired *t*-test *p* = 2.58 × 10⁻²⁹; Wilcoxon *p* = 7.79 × 10⁻¹⁰). However, median ALPS asymmetry remained largely unchanged (reference 8.73% versus pool-optimised 8.81%; paired *t*-test *p* = 0.195; Wilcoxon *p* = 0.369). Numerator asymmetry similarly showed no significant improvement, while denominator asymmetry decreased only modestly. These findings indicate that association ROI misalignment contributes to anatomical contamination but does not fully explain reference ALPS instability, motivating the subsequent multi-criteria optimisation framework.

### 3.3 Multi-criteria optimisation improves anatomical specificity and reduces hemispheric asymmetry

Building on the anatomical overlap optimisation stage, a multi-criteria optimisation framework was developed incorporating hemispheric ALPS asymmetry, left–right bias, component stability, and directional fibre purity (Figure 2A). Candidate ROI configurations were evaluated within anatomically constrained search regions and ranked using a weighted composite scoring framework.

**Figure 2.**
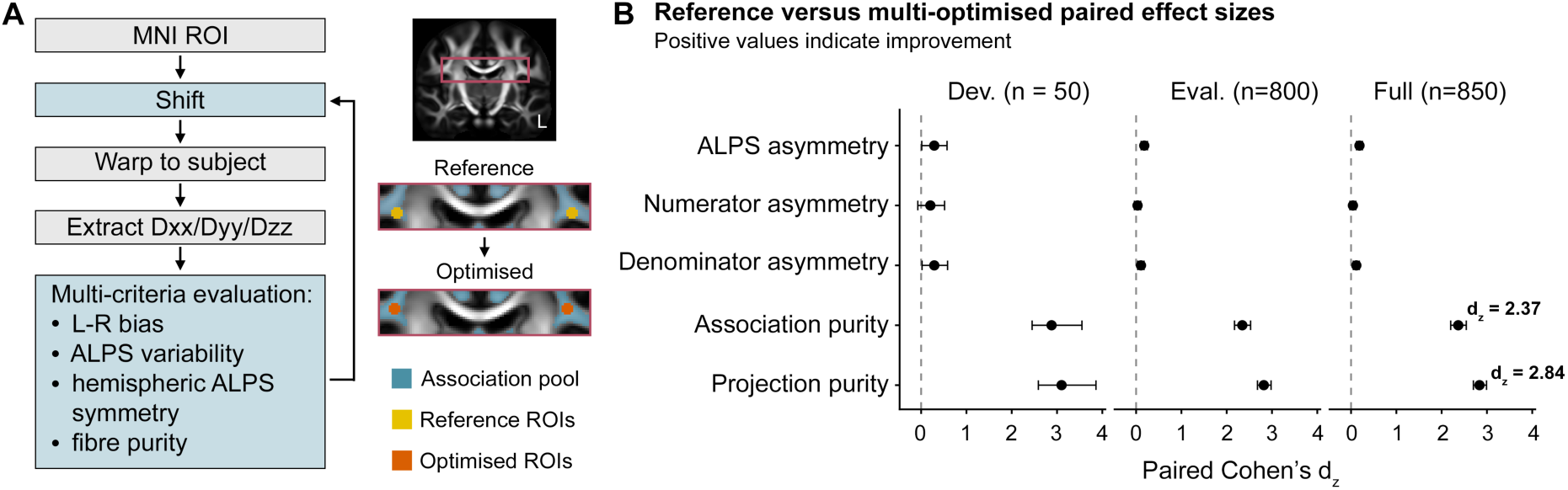
Multi-criteria optimisation improves anatomical specificity and hemispheric consistency of DTI-ALPS measurements. **(A)** Overview of the optimisation framework. Candidate ROI locations were generated within anatomically constrained fibre pools and evaluated using a multi-criteria scoring framework incorporating left–right bias, ALPS variability, hemispheric symmetry, and directional fibre purity. Reference ROIs (yellow) were shifted toward the tract core to generate optimised ROIs (orange). **(B)** Reference versus multi-optimised paired effect sizes across development (n = 50), independent evaluation (n = 800), and combined (n = 850) cohorts. Positive values indicate improvement following optimisation. Optimisation produced modest but consistent reductions in ALPS, numerator, and denominator asymmetry, while generating large improvements in association and projection fibre purity. Error bars indicate 95% confidence intervals for paired Cohen’s *d*_z_. Values shown for the full cohort are association purity (*d*_z_ = 2.37) and projection purity (*d*_z_ = 2.84).

Multi-criteria optimisation identified a final ROI configuration comprising projection ROIs centred at (26, −14, 22) mm and (−28, −12, 22) mm and association ROIs centred at (37, −13, 28) mm and (−38, −14, 28) mm in MNI space for the left and right hemispheres, respectively. These coordinates were subsequently fixed and applied without modification to the independent HCP-YA evaluation (n = 800) and full (n = 850) HCP-YA datasets. For the HCP-HA cohort, pool optimisation and multi-criteria optimisation were repeated independently using the 150 HCP-HA participants, allowing age-cohort-specific optimised ROI solutions to be derived and compared with reference measurements.

Application of the final optimised ROI configuration to the independent evaluation cohort (n = 800) demonstrated significant reductions in hemispheric ALPS asymmetry relative to reference ROIs. Median ALPS asymmetry decreased from 12.85% to 11.95% (paired *t*-test *p* = 4.36 × 10⁻⁷; Wilcoxon *p* = 1.47 × 10⁻⁶) (Figure 2B). Although the absolute reduction was modest (−0.9 percentage points), the improvement was consistently observed across the independent evaluation cohort and was not derived from the optimisation dataset itself.

Improvements in ALPS stability were accompanied by substantial increases in directional specificity. Association fibre purity increased from a median of 1.89 to 2.57, while projection fibre purity increased from 2.15 to 2.48 in the evaluation cohort. Both effects were highly significant (*p* < 10⁻¹³²) and corresponded to very large paired effect sizes (association purity *d*_z_ = 2.34; projection purity *d*_z_ = 2.82). Similar effects were observed in the development cohort and in the combined 850-subject dataset (Figure 2B).

Notably, optimisation produced only minor changes in bilateral ALPS magnitude itself, indicating that the principal effect of optimisation was not to alter ALPS values but rather to improve anatomical specificity and reduce hemispheric instability. Collectively, these findings demonstrate that reference ALPS variability arises in part from anatomically suboptimal ROI placement and that a multi-criteria optimisation strategy can substantially improve directional purity while providing measurable improvements in hemispheric consistency.

Association fibre-pool optimisation produced large improvements in anatomical specificity, and the final multi-criteria solution remained close to the pool-optimised coordinates. However, improvements in ALPS stability only emerged after incorporation of projection ROI optimisation and component-level stability constraints, indicating that reference ALPS variability reflects both association ROI misplacement and broader geometric instability of the underlying tensor components.

### 3.4 HCP-YA covariate analysis motivates ageing-cohort validation

To determine whether anatomical and demographic covariates contributed to DTI-ALPS variability even in young adulthood, multivariable models were fitted in the full HCP-YA cohort with age, sex and lateral ventricular volume as predictors. In this restricted young adult cohort, age explained little variance in either reference or multi-optimised ALPS metrics after accounting for sex and ventricular volume.

By contrast, lateral ventricular volume was consistently associated with bilateral ALPS and directional purity measures. For reference ALPS, larger ventricular volume was associated with lower bilateral ALPS (β = −0.138, FDR-adjusted *p* = 3.14 × 10⁻⁴), while for multi-criteria optimised ALPS this association was stronger (β = −0.343, FDR-adjusted *p* = 8.98 × 10⁻²⁴). Multivariable model fit likewise increased from R² = 0.046 for reference ALPS to R² = 0.165 for multi-criteria optimised ALPS. Ventricular volume was also strongly associated with projection purity for both reference and multi-criteria optimised measurements (β ≈ 0.38, FDR-adjusted *p* < 2 × 10⁻²⁷).

These findings indicate that even in young adults, ALPS is influenced by local anatomy surrounding the ventricles, with ventricular volume emerging as a stronger covariate than age. Because the HCP-YA cohort has a restricted age range and limited age-related ventricular enlargement, we next examined whether these relationships were amplified in an independent healthy ageing cohort spanning a broader range of ventricular morphology.

### 3.5 Optimisation generalises to an independent healthy ageing cohort

To determine whether the optimisation framework remained effective in the presence of age-related anatomical variation, pool optimisation and multi-criteria optimisation were repeated independently in the HCP-HA cohort (n = 150).

Interestingly, reference ALPS asymmetry was lower in HCP-HA (8.35%) than in HCP-YA (12.85%), indicating that the reference coordinate system performed comparatively better in the ageing cohort despite greater anatomical variability.

Association fibre-pool optimisation identified anatomically improved association ROI locations centred at (38, −12, 27) mm and (−37, −15, 27) mm for the left and right hemispheres, respectively. Independent optimisation identified remarkably similar anatomical solutions with the final ROI coordinates differing by only 1–2 mm from those identified in the HCP-YA cohort. Relative to the reference coordinates, optimisation increased median fibre-pool overlap from 82.6% to 100% for the left association ROI and from 91.3% to 100% for the right association ROI (both Wilcoxon *p* < 10⁻¹⁹), confirming that the reference association ROIs were partially displaced from the highest-density association fibre core in the ageing cohort.

Subsequent multi-criteria optimisation identified a final ROI configuration comprising projection ROIs centred at (26, −14, 22) mm and (−26, −12, 24) mm and association ROIs centred at (38, −11, 27) mm and (−37, −14, 28) mm in MNI space for the left and right hemispheres, respectively. These locations were selected using the same weighted optimisation framework applied in the HCP-YA cohort and again, differed by only 1–2 mm (Table 2).

**Table 2.**
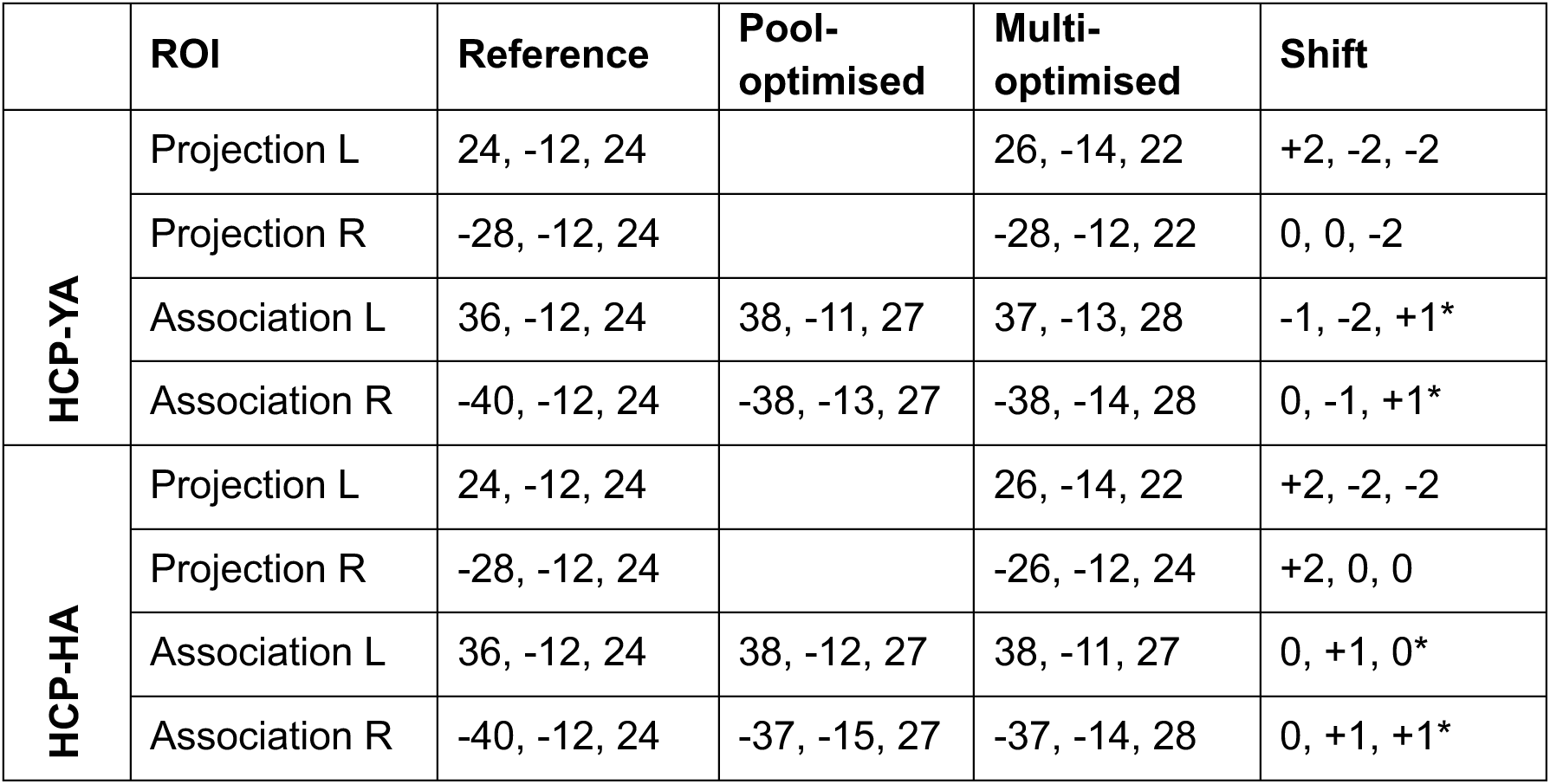
Final MNI coordinates following pool and multi-criteria optimisation. Reference coordinates were derived from published DTI-ALPS implementations. Pool-optimised coordinates were obtained by maximising overlap with directional fibre pools, whereas multi-optimised coordinates were derived using the multi-criteria optimisation framework. Shifts are reported in millimetres relative to the preceding optimisation stage; association ROI shifts (*) are given relative to the pool-optimised coordinates. Coordinates are reported in MNI space.

|  | ROI | Reference | Pool-optimised | Multi-optimised | Shift |
| --- | --- | --- | --- | --- | --- |
| HCP-YA | Projection L | 24, -12, 24 |  | 26, -14, 22 | +2, -2, -2 |
|  | Projection R | -28, -12, 24 |  | -28, -12, 22 | 0, 0, -2 |
|  | Association L | 36, -12, 24 | 38, -11, 27 | 37, -13, 28 | -1, -2, +1* |
|  | Association R | -40, -12, 24 | -38, -13, 27 | -38, -14, 28 | 0, -1, +1* |
| HCP-HA | Projection L | 24, -12, 24 |  | 26, -14, 22 | +2, -2, -2 |
|  | Projection R | -28, -12, 24 |  | -26, -12, 24 | +2, 0, 0 |
|  | Association L | 36, -12, 24 | 38, -12, 27 | 38, -11, 27 | 0, +1, 0* |
|  | Association R | -40, -12, 24 | -37, -15, 27 | -37, -14, 28 | 0, +1, +1* |

Comparison of reference and multi-optimised ALPS measurements demonstrated significant improvements in hemispheric consistency. Median ALPS asymmetry decreased from 8.35% to 5.14% (paired *t*-test *p* = 1.92 × 10⁻⁶; Wilcoxon *p* = 3.58 × 10⁻⁶), corresponding to a 32.0% relative reduction and a moderate paired effect size (*d*_z_ = 0.40). Denominator asymmetry also decreased from 7.83% to 6.02% (*p* = 1.32 × 10⁻³), whereas numerator asymmetry was largely unchanged.

As observed in the HCP-YA cohort, optimisation produced substantially larger improvements in directional specificity than in ALPS asymmetry itself. Median association purity increased from 1.85 to 2.61 (+40.9%; *d*_z_ = 3.10), while projection purity increased from 2.12 to 2.42 (+15.3%; *d*_z_ = 2.75). Both effects were highly significant (paired *t*-tests *p* < 10⁻⁷⁰). Collectively, these findings demonstrate that the optimisation framework generalises to an independent ageing population and consistently improves anatomical specificity while reducing hemispheric variability.

### 3.6 Ventricular anatomy explains substantially more ALPS variance than chronological age

To determine whether ALPS variability was more strongly related to ageing itself or to age-associated anatomical changes, multivariable models incorporating age, sex, and ventricular volume were evaluated in the HCP-HA cohort.

For bilateral ALPS, ventricular volume consistently explained substantially more variance than chronological age. In reference ALPS measurements, ventricular–volume-only models explained 41.3% of variance compared with 20.3% for age-only models. Following optimisation, explanatory power increased further, with ventricular volume alone explaining 58.1% of variance compared with 37.5% for age alone. Full models incorporating age, sex, and ventricular volume explained 41.8% and 60.5% of variance for reference and multi-optimised ALPS, respectively.

Partial F-tests demonstrated that ventricular volume contributed substantial independent explanatory power beyond age and sex. For reference ALPS, ventricular volume increased model fit by ΔR² = 0.210 (*p* = 2.11 × 10⁻¹¹), whereas age contributed no additional variance after accounting for ventricular volume and sex (ΔR² < 0.001, *p* = 0.894). Similar results were observed for multi-optimised ALPS, where ventricular volume increased explanatory power by ΔR² = 0.228 (*p* = 4.02 × 10⁻¹⁶), while age again contributed little independent variance (ΔR² = 0.008, *p* = 0.084).

Analysis of unique variance components confirmed these findings. In the full model, ventricular volume accounted for the majority of explainable variance in bilateral ALPS, whereas age and sex contributed comparatively little additional information. Optimisation increased the proportion of variance attributable to ventricular anatomy, indicating that the optimised ALPS metric is more strongly coupled to underlying periventricular morphology than the reference measurement.

Non-linear modelling provided only modest improvements over linear fits. Natural spline models increased explained variance by ΔR² = 0.058 for reference ALPS (*p* = 4.92 × 10⁻⁴) and by ΔR² = 0.016 for multi-optimised ALPS (*p* = 0.050), suggesting that the dominant relationship between ALPS and ventricular anatomy is well approximated by a linear model.

Collectively, these findings indicate that ventricular anatomy, rather than chronological age itself, is the principal determinant of ALPS variability in healthy ageing. This observation motivated subsequent investigation of periventricular tissue deformation using DBM and log-Jacobian analyses.

### 3.7 ALPS is associated with local periventricular tissue deformation

To investigate the anatomical basis of the ventricular volume association, DBM analyses were performed in the HCP-HA cohort using voxel-wise log-Jacobian maps derived from T1-weighted structural MRI.

For both reference and multi-optimised ALPS measurements, lower bilateral ALPS values were associated with significantly greater ventricular and periventricular expansion (TFCE-corrected *p* < 0.05; Figure 3A). Significant clusters were concentrated around the lateral ventricles, including the frontal horns, ventricular body, and temporal horn regions, indicating that ALPS reductions were specifically linked to local ventricular enlargement rather than diffuse global brain atrophy.

**Figure 3.**
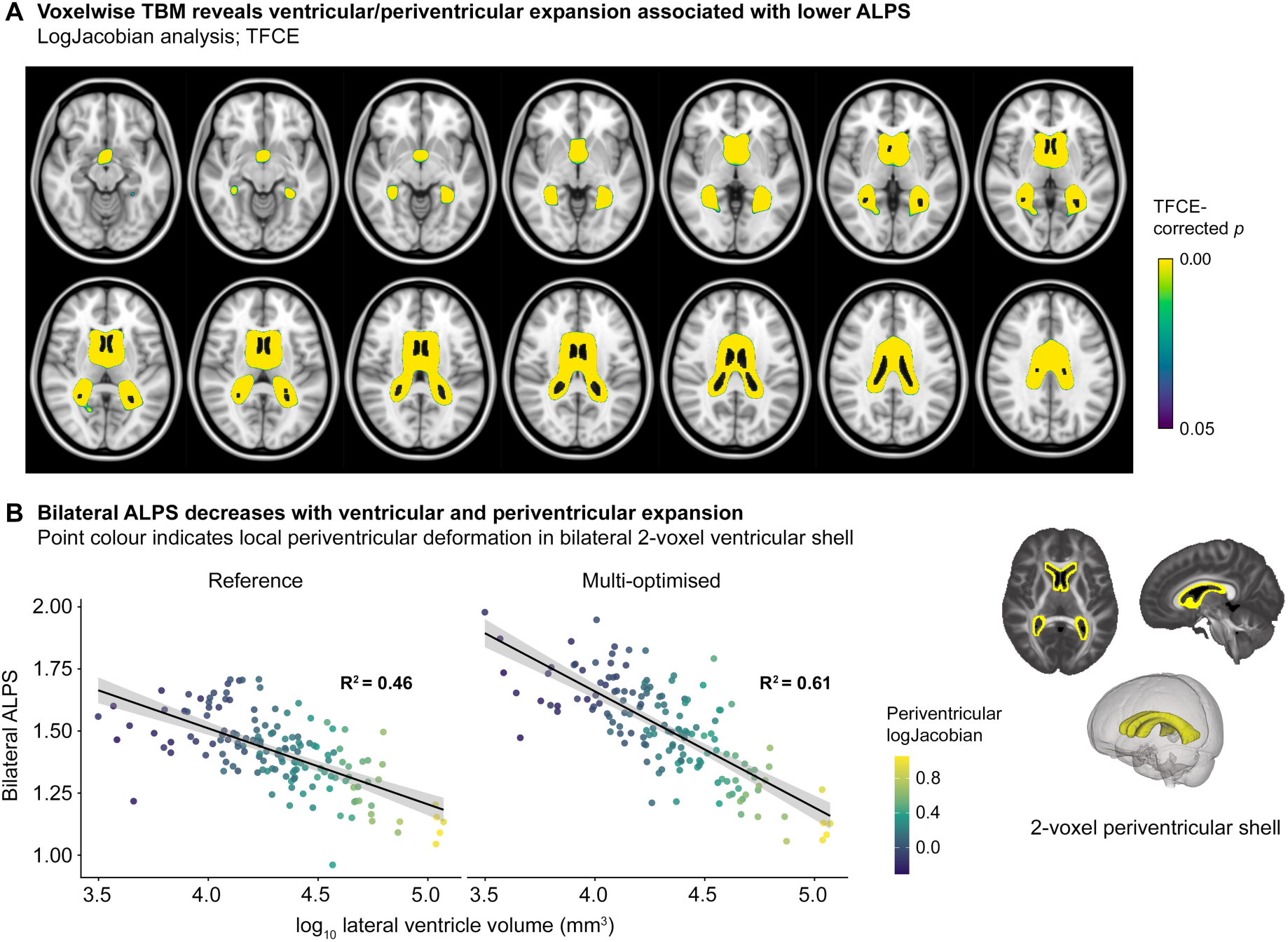
Lower ALPS values are associated with ventricular enlargement and periventricular tissue expansion. **(A)** Voxel-wise deformation-based morphometry (DBM) analysis relating bilateral ALPS to log-Jacobian values in the HCP-HA cohort. Significant TFCE-corrected clusters (*p* < 0.05) indicate regions where lower ALPS values are associated with greater local tissue expansion. Effects are concentrated around the lateral ventricles and adjacent periventricular white matter. **(B)** Relationship between bilateral ALPS and ventricular anatomy. Scatter plots show bilateral ALPS versus lateral ventricular volume for reference and multi-optimised ROIs. Point colour indicates mean log-Jacobian within an independently defined bilateral 2-voxel periventricular shell surrounding the lateral ventricles. Optimisation strengthened the association between ALPS and ventricular morphology (*R*² = 0.46 reference; *R*² = 0.61 multi-optimised). Insets show the anatomical definition of the 2-voxel periventricular shell used for deformation analyses.

To quantify this relationship, mean log-Jacobian values were extracted from an *a priori* defined bilateral 2-voxel periventricular shell surrounding the lateral ventricles (Figure 3B). Strong inverse relationships were observed between bilateral ALPS and ventricular volume for both reference and multi-optimised measurements, with optimisation strengthening the anatomical association. Reference ALPS explained 46.4% of variance in the periventricular deformation model (R² = 0.464), whereas multi-optimised ALPS explained 61.4% (R² = 0.614).

Multivariable analyses incorporating age, sex, ventricular volume and shell log-Jacobian demonstrated that local periventricular deformation contributed unique explanatory power beyond ventricular volume alone for reference ALPS measurements. In the reference model, the 2-voxel shell explained 4.6% unique variance, whereas ventricular volume contributed <0.1% unique variance after accounting for shell deformation. Partial F-tests confirmed that shell log-Jacobian significantly improved model fit beyond age, sex and ventricular volume (ΔR² = 0.046, FDR-corrected *p* = 0.003), whereas ventricular volume did not contribute additional explanatory power once shell deformation was included (FDR-corrected *p* = 0.725).

In contrast, for multi-optimised ALPS measurements, the relationship was dominated by ventricular volume itself. The 2-voxel shell explained only 0.8% unique variance, and shell deformation no longer significantly improved model fit after accounting for ventricular volume (FDR-corrected *p* = 0.176). Together, these findings suggest that optimisation increased coupling between ALPS and gross ventricular morphology while reducing sensitivity to localised periventricular deformation effects.

## 4. DISCUSSION

In this study, I systematically evaluated the effects of ROI positioning, ventricular morphology, and periventricular deformation on DTI-ALPS using large-scale diffusion MRI data from the HCP-YA and HCP-HA cohorts (Van Essen, Ugurbil, et al., 2012). Employing a multi-stage optimisation strategy incorporating fibre-pool alignment, hemispheric symmetry, component stability, and directional purity, I identified ROI configurations that substantially improved measurement performance in young adults. However, I found that optimisation did not eliminate hemispheric differences entirely. Further, ALPS demonstrated strong associations with ventricular anatomy and periventricular deformation, suggesting that ventricular morphology is a major anatomical determinant of ALPS variability. Collectively, these findings indicate that DTI-ALPS is shaped by a combination of methodological implementation and underlying neuroanatomy, with important implications for interpretation of the metric as a biomarker of putative glymphatic function.

A principal finding was the unexpectedly large degree of hemispheric variability present in reference ALPS measurements. Across 850 healthy young adults, reference ROIs produced a median left-right ALPS difference of approximately 12.5%, with nearly one-third of subjects exhibiting hemispheric differences exceeding 20%. Importantly, similar asymmetry was observed within the numerator and denominator components of the ALPS ratio, indicating that substantial instability exists in the underlying tensor measurements themselves. These findings suggest that apparent bilateral agreement in ALPS may partially reflect cancellation of opposing component-level asymmetries rather than true measurement stability. Such variability has important implications for studies interpreting relatively small group differences in ALPS as evidence of altered glymphatic function.

Anatomical optimisation substantially improved performance in the HCP-YA cohort. Pool optimisation corrected a systematic offset between reference association ROIs and underlying fibre anatomy, resulting in improved fibre overlap and reduced hemispheric asymmetry. Subsequent multi-criteria optimisation further improved directional purity and component stability while maintaining anatomical plausibility. Notably, optimisation effects were highly reproducible, with similar improvements observed in both the development and independent validation cohorts. Together, these findings demonstrate that ROI placement represents a major source of avoidable measurement variability. Importantly, the largest improvements were observed in directional fibre purity rather than ALPS magnitude itself, indicating that optimisation primarily improves anatomical validity and measurement robustness rather than altering the biological signal being sampled.

Interestingly, optimisation solutions were remarkably similar across cohorts, differing by only 1–2 mm despite substantial differences in age and ventricular morphology. Nevertheless, the optimal ROI locations were not identical, indicating that subtle age-related anatomical changes influence the location of maximal fibre specificity. These findings argue against the existence of a universally optimal ALPS ROI while demonstrating that the optimisation framework itself is robust across independent populations.

Notably, optimisation did not eliminate asymmetry entirely. Residual hemispheric differences persisted even under the final multi-optimised configuration, suggesting the presence of a lower bound of variability that cannot be removed through ROI refinement alone. This observation is consistent with the known anatomical complexity of periventricular white matter, where crossing fibres, fibre dispersion, local curvature, and hemispheric structural differences may all influence directional diffusivity measurements (Schilling et al., 2025). The persistence of asymmetry despite extensive optimisation suggests that a component of ALPS variability reflects genuine anatomical heterogeneity rather than methodological error.

The DTI-ALPS literature has adopted several approaches to hemispheric sampling, including unilateral left-sided ROIs (Taoka et al., 2017; Yun et al., 2025), bilateral averaging (Han et al., 2023; S. Liu et al., 2024; X. Liu et al., 2024; Satpathi et al., 2025; Sharkey et al., 2024; Yu et al., 2026), and separate hemispheric analyses (Ayral et al., 2025; Georgiopoulos et al., 2024; Qiu et al., 2025; Roura et al., 2025; Taoka et al., 2022; Yu et al., 2026). The original ALPS implementation used left-sided ROIs based on the larger calibre of dominant-hemisphere association fibres and the perceived stability of ROI placement (Taoka et al., 2017). While recent studies frequently average left and right measurements to reduce sensitivity to head orientation and local anatomical variation, the magnitude of hemispheric variability has not been systematically quantified. The present findings provide quantitative evidence that hemispheric variability is considerably larger than has generally been appreciated. The observation that numerator and denominator components exhibit substantially greater asymmetry than the final ALPS ratio further suggests that bilateral averaging may partially mask instability in the underlying tensor measurements. Future studies should therefore report hemispheric measurements separately where possible and explicitly consider hemispheric variability when interpreting small between-group differences.

The present findings complement recent work questioning the biological specificity of ALPS. Schilling and colleagues demonstrated that ALPS measurements are strongly influenced by white matter geometry, fibre crossings, axonal undulations, and vascular orientation assumptions, raising important concerns regarding interpretation of ALPS as a direct measure of glymphatic function (Schilling et al., 2025). Similarly, Taoka and colleagues have emphasised that the ALPS index should be interpreted as a measure of directional water diffusivity at the level of the lateral ventricles rather than a direct surrogate of glymphatic transport (Taoka et al., 2024). The current study extends these observations by demonstrating that ROI definition and ventricular anatomy constitute additional major determinants of ALPS variability. Together, these findings suggest that ALPS reflects a complex interaction between tissue microstructure, ventricular morphology, and methodological implementation, requiring careful interpretation when used as a putative biomarker of glymphatic function.

The HCP-HA cohort provided further evidence that ventricular anatomy exerts a substantial influence on ALPS measurements. Across multiple regression models, ventricular volume emerged as the strongest and most consistent predictor of ALPS, whereas chronological age contributed relatively little additional explanatory value once ventricular size was considered. These findings suggest that previously reported age-related reductions in ALPS (Han et al., 2023; Roura et al., 2025; Satpathi et al., 2025) may, at least in part, reflect age-associated ventricular enlargement rather than ageing itself. Consistent with this interpretation, ventricular volume explained a substantially greater proportion of ALPS variance than age. This distinction is important because ventricular enlargement is only one component of the ageing process and varies considerably between individuals. The present findings therefore suggest that anatomical variation surrounding the lateral ventricles, rather than ageing *per se*, is a principal contributor to ALPS variability.

DBM provided complementary evidence linking ALPS to ventricular anatomy. Lower ALPS values were associated with widespread ventricular and periventricular expansion, with effects localised predominantly to tissue immediately surrounding the lateral ventricles. Notably, these regions closely mirrored the spatial pattern of age-related ventricular enlargement, suggesting that ALPS is sensitive not only to ventricular size but also to deformation of the surrounding white matter architecture. Because the ALPS ROIs are positioned immediately adjacent to the lateral ventricles, these findings provide a plausible anatomical explanation for the observed ventricular-volume effects. Small changes in ventricular morphology are likely to alter the surrounding fibre geometry, local tissue deformation, and relative orientation of perivascular spaces, each of which has the potential to influence directional diffusivity measurements.

These findings are consistent with observations in idiopathic normal pressure hydrocephalus, where Evans index emerged as one of the strongest predictors of ALPS and ventriculomegaly showed a robust inverse relationship with ALPS values (Georgiopoulos et al., 2024). Georgiopoulos et al. interpreted this association as evidence that ventricular enlargement may impair glymphatic exchange through altered CSF dynamics. While the present findings do not exclude this possibility, they further suggest that structural reorganisation of periventricular tissue may contribute substantially to the observed relationship. Taken together, the covariate and DBM analyses indicate that ALPS is closely coupled to ventricular morphology and surrounding tissue deformation, raising the possibility that ALPS functions, at least in part, as a marker of periventricular structural organisation rather than a measure of glymphatic transport alone.

Recent work has questioned whether ALPS primarily reflects perivascular diffusivity, demonstrating that widespread radial tensor asymmetry (λ_2_ > λ_3_) exists throughout white matter and decreases with ageing and cognitive impairment (Wright et al., 2024). Wright et al. proposed that intrinsic white matter microstructure may therefore contribute substantially to ALPS measurements. The present study identifies a complementary source of variability, demonstrating that ROI placement, fibre-pool alignment, ventricular morphology, and hemispheric asymmetry also exert substantial effects on ALPS quantification. Together, these findings suggest that ALPS should be interpreted as a composite anatomical diffusion biomarker influenced by both tissue microstructure and measurement geometry rather than a direct or isolated measure of glymphatic function.

Importantly, these findings should not be interpreted as evidence that ALPS lacks biological relevance. Rather, they demonstrate that ALPS is influenced by multiple interacting anatomical and methodological factors that require consideration when interpreting the metric. Accounting for ROI definition and ventricular morphology is therefore likely to improve, rather than diminish, the utility of ALPS as a neuroimaging biomarker by reducing measurement variability and improving biological specificity.

Several limitations should be acknowledged. First, no universally accepted DTI-ALPS ROI definition currently exists (Taoka et al., 2024). Although the present study adopted a commonly used literature-derived configuration as the reference implementation, alternative ROI coordinates and geometries have been reported, further highlighting the need for standardisation of ALPS methodology. Second, optimisation was performed using tensor-derived directional information rather than direct histological or vascular validation. While fibre-pool alignment provides an objective anatomical framework, it cannot establish whether optimised ROIs better reflect perivascular fluid transport. Likewise, although ventricular morphology explained a substantial proportion of ALPS variance, the cross-sectional design cannot determine whether ventricular enlargement directly influences ALPS or whether both reflect common underlying biological processes. Finally, although large HCP cohorts provide excellent statistical power and image quality, findings may not generalise directly to clinical populations or datasets acquired with different imaging protocols.

In conclusion, ROI definition represents a major source of variability in DTI-ALPS measurements, and anatomically informed optimisation substantially improves measurement stability in healthy young adults. However, a residual asymmetry floor remains even after optimisation, suggesting that part of ALPS variability reflects intrinsic anatomical complexity. More importantly, ventricular morphology and periventricular tissue deformation emerged as dominant determinants of ALPS variation, exceeding the contribution of chronological age alone. These findings provide an anatomical framework for interpreting DTI-ALPS measurements and support a shift from viewing ALPS as a direct surrogate of glymphatic function toward understanding it as a composite diffusion biomarker shaped by both neuroanatomy and methodological implementation. Future studies should explicitly account for ventricular morphology and ROI definition when interpreting ALPS and consider these factors as integral components of study design rather than *post hoc* confounders. More broadly, the present findings emphasise the importance of rigorous anatomical validation when developing and applying diffusion MRI biomarkers intended to reflect underlying physiological processes.

## 5. Data and Code Availability

Human Connectome Project Young Adult and Healthy Aging data are available to qualified researchers through the Human Connectome Project under the applicable data use agreements. The custom analysis code supporting this study will be deposited in a public GitHub repository and archived with a DOI (e.g. Zenodo) upon acceptance of the manuscript.

## 6. Author Contributions

D.K.W. conceived and designed the study, developed the methodology, performed all analyses, interpreted the results, prepared the figures, and wrote the manuscript.

## 7. Funding

D.K.W. acknowledges the financial support of FightMND through a Mid-Career Research Fellowship (MCR-202503-01826_Wright).

## 8. Declaration of Competing Interests

D.K.W. has no conflict of interest.

## 9. Acknowledgements

HCP-Young adult data were provided by the Human Connectome Project, WU-Minn Consortium (Principal Investigators: David Van Essen and Kamil Ugurbil; 1U54MH091657) funded by the 16 NIH Institutes and Centers that support the NIH Blueprint for Neuroscience Research; and by the McDonnell Center for Systems Neuroscience at Washington University. HCP-Aging adult brain connectome (AABC) release 2 data, methods used, and/or research reported in this publication were provided in whole or in part by the Aging Adult Vulnerability and Resiliency in the Aging Adult Brain Connectome (AABC) project (U19AG073585) and the Human Connectome Project in Aging (HCP-A, U01AG052564) funded by the National Institute of Aging of the National Institutes of Health. HCP-A was further supported by funds provided by the McDonnell Center for Neuroscience at Washington University in St. Louis.

The content is solely the responsibility of the author and does not necessarily represent the official views of the National Institutes of Health.

D.K.W. used ChatGPT (OpenAI, San Francisco, CA, USA) as a research support tool during study design, statistical analysis, code development, figure preparation, literature review, and manuscript drafting. All analytical workflows, results, interpretations, and final manuscript content were independently verified and approved by the author.

## Notes

### Competing Interest Statement

The authors have declared no competing interest.

